# Modeling presynaptic inhibition by the amyloid precursor protein demonstrates one potential mechanism for preventing runaway synaptic modification in Alzheimer’s disease

**DOI:** 10.1101/2025.08.11.669740

**Authors:** Dylan Barber, Michael E. Hasselmo, Heather C. Rice

## Abstract

**INTRODUCTION:** Previous simulations of Hebbian associative memory models inspired the malignant synaptic growth hypothesis of Alzheimer’s disease (AD), which suggests that cognitive impairments arise due to runaway synaptic modification resulting from poor separation between encoding and retrieval.

**METHODS:** We computationally model presynaptic inhibition by the recently identified interaction of soluble amyloid precursor protein (sAPPα) with the γ-aminobutyric acid type B receptor (GABA_B_R) as one potential biological mechanism which can enhance separation between encoding and retrieval.

**RESULTS:** Simulations predict that the dual effect of sAPPα on long-term potentiation and presynaptic inhibition of glutamatergic synapses maintains effective associative memory function and prevents runaway synaptic modification. Moreover, computational modeling predicts that sAPPα, which interacts with the 1a isoform of GABA_B_R, is more effective at stabilizing associative memory than the GABA_B_R agonist Baclofen.

**DISCUSSION:** Molecular mechanisms that enhance presynaptic inhibition, such as sAPPα-GABA_B_R1a signaling, are potential therapeutic targets for preventing cognitive impairments in AD.

## 1. BACKGROUND

Prominent hypotheses of Alzheimer’s disease (AD) center on molecular pathways such as amyloid and tau. However, the malignant synaptic growth hypothesis [1–3] proposes that AD arises from an imbalance of synaptic strengthening leading to runaway synaptic modification (malignant synaptic growth) that exacerbates amyloid and tau pathologies. The imbalance could arise from variations in molecular mechanisms for associative memory function in cortical structures [4–6] that regulate the separation of encoding and retrieval.

Here, encoding refers to the NMDA-dependent process of synaptic modification to store novel memories, and retrieval refers to the spread of activity via glutamate release at modified synapses to recall previously stored memories.

Malignant synaptic growth is typically prevented in associative memory models such as Hopfield networks [7–11] by complete separation of encoding and retrieval [7–11]. However, because glutamate release is necessary for both memory retrieval and encoding via NMDA-dependent synaptic modification, the overlap of encoding and retrieval is inevitable in biological circuits. Mathematical models presented here show how variation of molecular pathways that regulate the separation of encoding and retrieval can allow malignant synaptic growth that increases both the total number of synapses, and the total strength of individual synapses [1, 12–14]., These models can account for the initial appearance of AD tangle pathology in the hippocampus and entorhinal cortex [15, 16], the dysfunction of synapses in the hippocampus [17, 18], the evidence of fMRI hyperactivation in the hippocampus in early AD [19–22]. [19, 20, 23], and spread of disease pathology along pathways associated with memory consolidation [24].

A number of molecular mechanisms can mediate separation of encoding and retrieval through presynaptic inhibition, including activation of muscarinic acetylcholine receptors [25], GABAB receptors [26] and mGluR receptors. The balance can also be influenced by A*β* regulation of synaptic plasticity. Moreover, runaway synaptic transmission can be prevented by mechanisms of homeostatic plasticity, including physical synapse remodeling, presynaptic regulation of release mechanisms, regulation of synaptic AMPA and NMDA receptors, and GPCRs (including GABA_B_R) [27–34]. This paper focuses on sAPPα-GABA_B_R1a signaling [35, 36] as one mechanism which might contribute to prevention of runaway synaptic modification (RSM) (Figure 1). This consideration is particularly critical, as APP-targeting siRNA therapeutics are currently undergoing clinical evaluation for the treatment of Alzheimer’s disease and cerebral amyloid angiopathy (CAA) [37].

**Figure 1.**
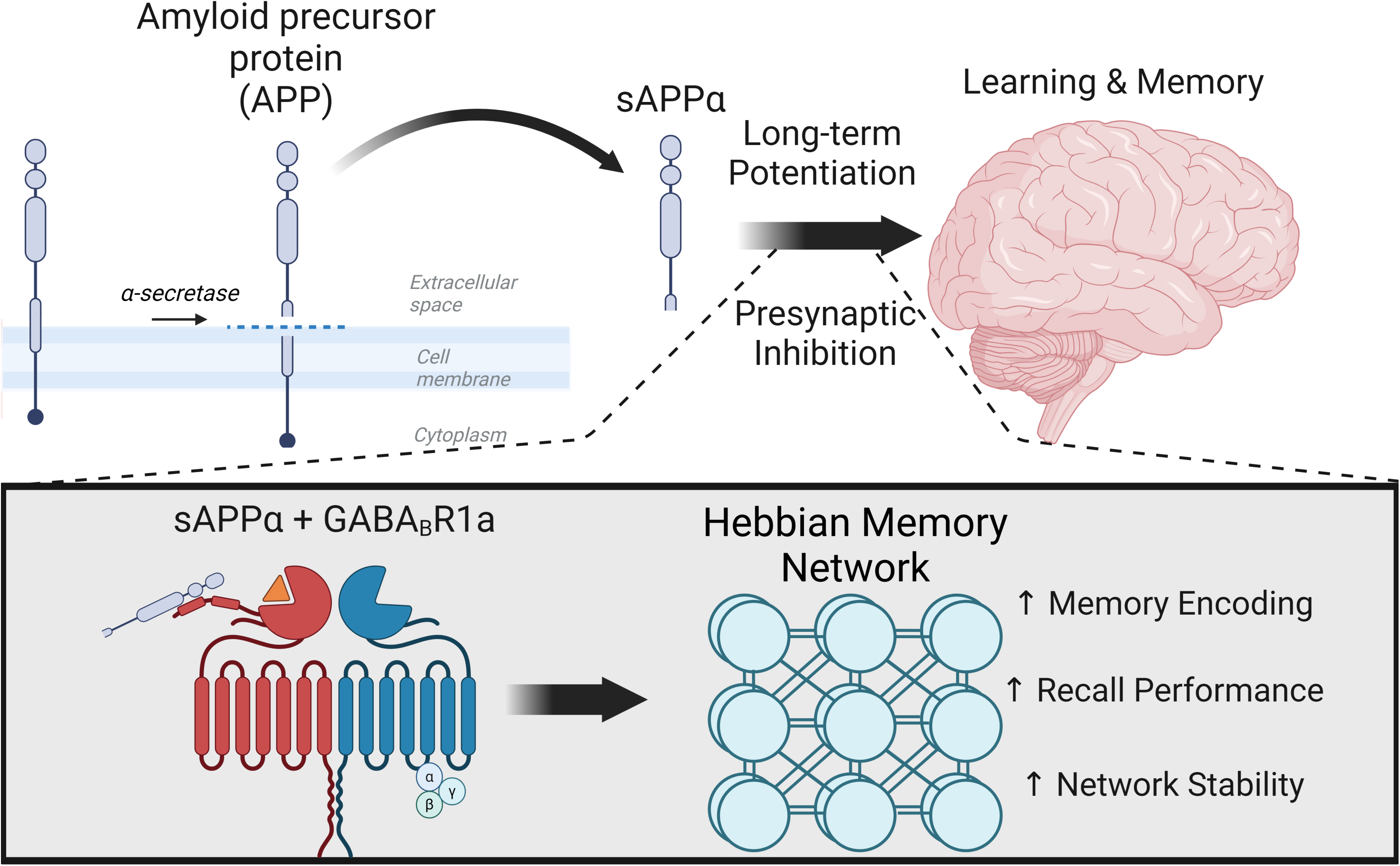
Proposed model of the protective role of sAPP signaling on brain networks and memory function. Our model proposes that an increase of sAPPα secretion stimulated by neuronal activity contributes to learning and memory. sAPPα increases both long term potentiation (LTP) and presynaptic inhibition (PreInhib). The GABA_B_ Receptor subunit 1A (GABA_B_R1A) mediates the effects of sAPP on presynaptic inhibition. Models of Hebbian associative learning indicate that this combined function of sAPP can increase network stability and increase memory recall performance by reducing interference between encoding and retrieval. Created in BioRender. Barber, D. (2025) https://BioRender.com/l41i127

APP is a type I transmembrane protein that undergoes sequential proteolytic processing to generate multiple peptides including Amyloid-β (Aβ), the primary constituent of amyloid plaques found in Alzheimer’s disease (AD) brains. The normal function of APP promotes learning [38–40] and regulates both encoding via enhancing long-term potentiation [41, 42] and retrieval via regulation of presynaptic inhibition [35]. The secreted APP ectodomains (sAPP), which are generated during the initial cleavage event by α- or β-secretases, appear to mediate these functions of APP, and could switch the network between encoding (when sAPP enhances LTP and causes presynaptic inhibition) and retrieval (when APP has not been cleaved).

Studies have addressed the functional role of sAPP*α*. In slices, sAPPα reduces synaptic activity and enhances long-term potentiation (LTP) of glutamatergic synapses [41, 43–45] potentially via nicotinic acetylcholine receptor 7 signaling and calcium permeable AMPA receptor recruitment [46, 47]. sAPPα also regulates dendritic spine density [46]. sAPPα rescues defects in LTP and spatial learning [46, 48, 49] in App knockout mice. The γ-aminobutyric acid type B receptor (GABA_B_R) has been shown to bind the APP ectodomain [35, 36] and mediate effects of sAPP on presynaptic inhibition of glutamate release [35, 36]. The specific mechanisms mediating the functional effects of this interaction are still under active investigation. These mechanisms may not stimulate canonical GABA_B_R signaling [50]; however, this is still being investigated. In these studies [35, 36, 50] the APP extracellular domain was found to specifically bind the 1a isoform of the GABA_B_R, whereas current GABA_B_R agonists, such as baclofen target both 1A and 1B isoforms.

We employed a standard Hebbian associative memory model to address the:

1. relationship between an increase in both long-term potentiation and presynaptic inhibition of glutamatergic synapses by sAPP
2. potential protective role of sAPP-GABA_B_R1a signaling on brain networks in Alzheimer’s disease
3. potential therapeutic implications of GABA_B_R1a-specific modulation as compared to non-isoform-specific modulation of GABA_B_R signaling
4. potential detrimental impacts of reducing APP expression by RNAi as a treatment for Alzheimer’s disease

## 2. METHODS

### 2.1 Small Circuit Example

#### 2.1.1 Associative memory model description of the physiological effects of sAPP-GABA_B_R1a binding

Memory models commonly focus on the formation of associations between different elements of a stored episodic memory (Fig. 2A). For example, one might form an association between a face and a place, for example when forming the episodic memory of meeting a new colleague at a reception. Standard models of associative memory link the pattern of neural activity p represented by a series of binary vectors in one population (i.e. the input layer) to a pattern of neural activity, also represented as binary vectors, r in another population (i.e. the output layer). Note that these binary vectors are not orthogonal (they contain overlapping active elements). The storage of correlated patterns has been known as an issue for the capacity of Hopfield networks [51, 52] that can be reduced by learning rules that decrease correlations, but here we focus on the breakdown of function enhanced by correlations.

**Figure 2.**
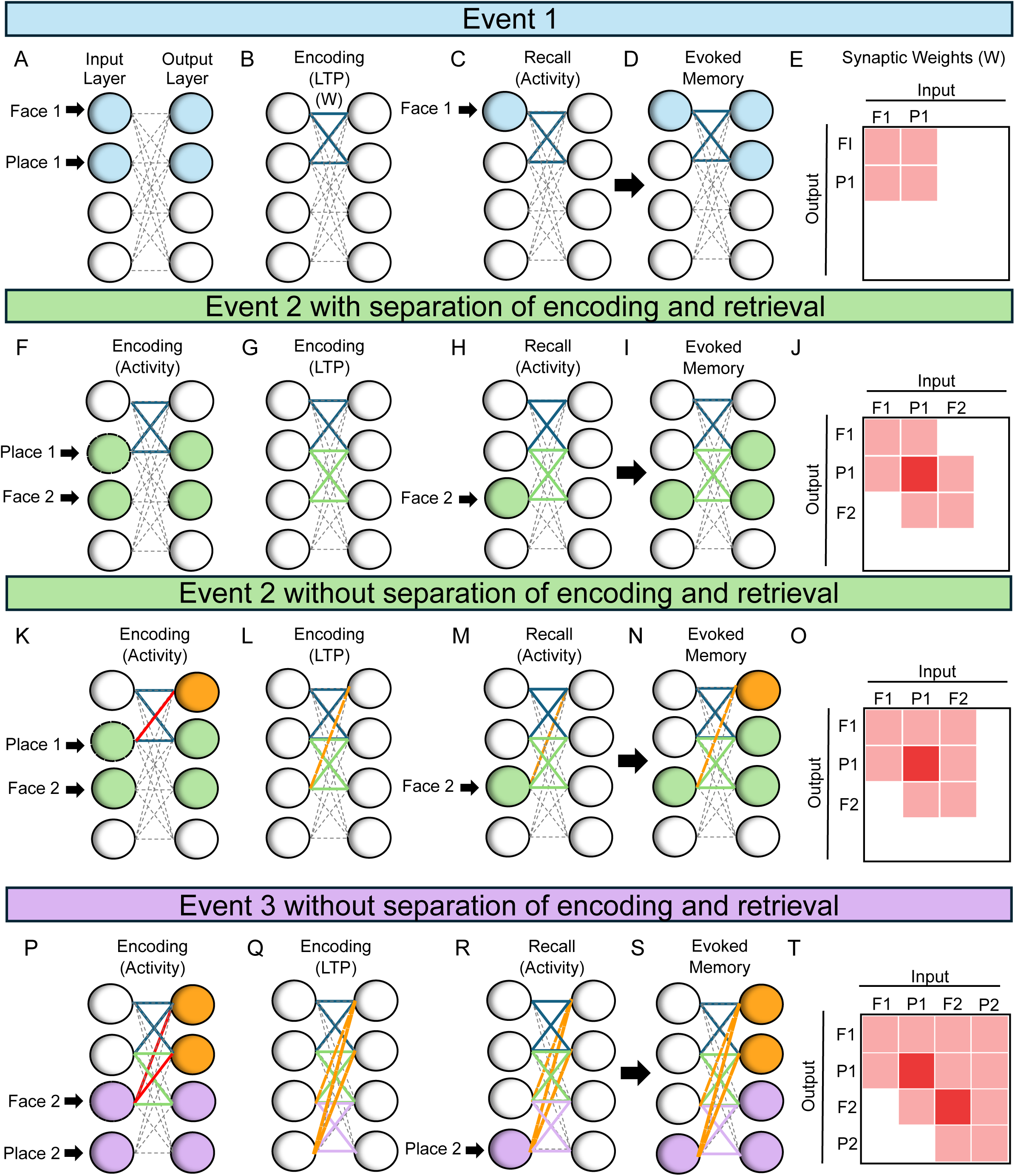
Models of Hebbian associative memory show effective function with separation of encoding and retrieval and undesired synaptic growth without this separation. A) In the first event, one population of activated neurons (left blue circles) responds to a sensory input representing a face and a place associated with the memory. Another output population (right blue circles) responds to the input in corresponding manner. B) During encoding, Hebbian synaptic modification (Encoding LTP) strengthens synapses between the activated neurons (thick blue lines). C) The input layer of neurons receives activity corresponding to the face 1 associated with place 1. This spreads across the matrix (blue lines). D) The input of face 1 evokes the full memory encoded earlier in step B in the output layer due to the recently modified synapses. E) The same pattern of strengthened synapses (blue lines) can be represented by pink shaded squares in a matrix of connections. F-J) With separation of encoding and retrieval, the input of a consecutive overlapping event 2 activates neurons (green circles) that have overlap of place 1 being associated with a novel face. F) With separation of encoding and retrieval, encoding activity only activates the neurons getting direct input (green circles). G) This encoding activity causes strengthening (LTP) of only the desired synapses (green lines) with no undesired strengthening. H) The input layer receives activity corresponding to face 2 that was associated with place 1. I) The full memory of face 2 and place 1 encoded from step G is recalled without interference. J) The 3 new strengthened synapses are shown as pink squares. K) Without separation between encoding and retrieval, encoding activity with retrieval involves neurons getting direct input (green circles) and spreading of retrieval activity across a previously strengthened synapse (red line and orange circle). L) This mixed postsynaptic activity causes strengthening (LTP) of not only the new association (thick green lines) but also additional undesired synapses are strengthened (orange dashed line). M) The input layer receives activity corresponding to face 2. N) The memory evoked by retrieval includes not only the correct memory (green) but also interference from activation of the postsynaptic neuron previously associated with face 1 (orange). O) The undesired synapse appears as a pink square in column F2, row F1. P) For event 3, without separation of encoding and retrieval, encoding activity with retrieval spreads to both the previous memories (red lines and orange circles). Q) This additional activity causes LTP of the new association (purple lines) and two undesired synapses being strengthened (orange dashed lines). R) The input layer receives activity corresponding to only place 2. S) The memory recalled in the output layer now includes even more interference from both face 1 and place 1 due to undesired modification from event 2. T) The map of the modified synapses now reflects the additional undesired synapses as pink squares in column P2, row F1, and column P2, row P1. This shows the undesired synaptic growth associated with interference from retrieval during encoding.

The associative memory is formed by a pattern of synapses W between the two populations. Figure 2A shows an example of associative memory connectivity for a single episode regarding a face and a place.

The mathematical description here first shows the breakdown in function using equations that are standard components of associative memory models with Hebbian modification [7–11]. The later equations then add parameters that reflect the stabilizing physiological effects of molecular pathways that regulate synaptic modification and synaptic transmission [1, 12, 13]. The equations effectively address the associative memory function used in most models of the hippocampal formation [4–6], though this associative memory model is very simple relative to the data showing detailed physiological differences between different anatomical subregions and sublayers of the hippocampus [53, 54]. More elaborate models of hippocampus have used ionic conductances to simulate spiking dynamics, but usually still rely on the same functional elements of associative memory with Hebbian synaptic modification for completion of memories [55] and Hebbian STDP for the replay of encoded sequences [56].

Most associative memory models encode new associations between patterns by using the Hebb rule, which causes a change in the strength of a matrix of synaptic weights W between neurons in the population (thick blue lines in Fig. 2B). This synaptic matrix is modified based on a vector *p* representing the array of activity in a first population of presynaptic neurons, and the vector *r* representing the array of activity across a second population of postsynaptic neurons. The Hebb rule for modification of synaptic weights results in the pattern of connectivity shown in Figure 2 B, and the matrix or map of connectivity shown in Figure 2E. Mathematically, this corresponds to the creation of this matrix from the outer product of the presynaptic pattern p and the postsynaptic pattern r, as represented by the following equation (where a vector transpose is indicated by T):

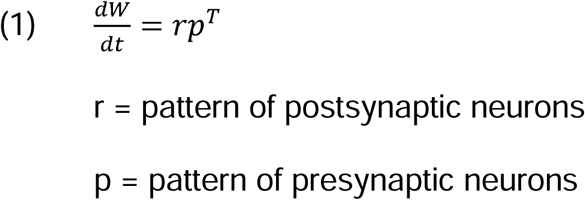

After encoding, the associative memory model can retrieve the association with corresponding input. A portion of the presynaptic activity pattern in the first population *p* is activated by sensory input from the outside world and acts as a cue for the memory. Additional memories can be encoded even with overlapping associations (Fig. 2F). For example, Figure 2F shows how input corresponding to the place encoded in the memory in Figure 2A can spread activity across the strengthened synapses (thick blue lines) to activate postsynaptic neurons that represent the retrieval of the memory previously encoded, but this retrieval is separated from the encoding step (Fig. 2G).

In the equations, this retrieval process is represented by the vector of activity *p* being multiplied by the matrix of synaptic weights W to generate a vector of postsynaptic activity r. Thus, the retrieval process is described by the equation:

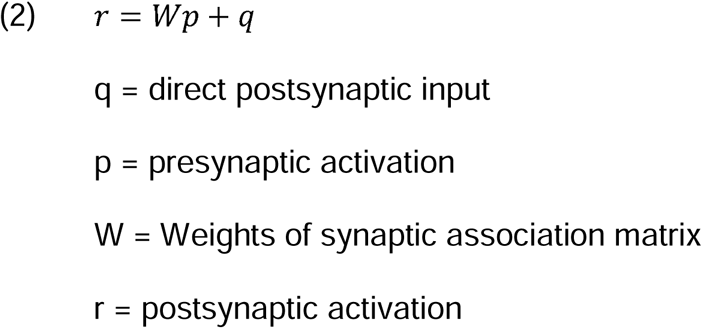

Where r is a vector representing the activity of all the neurons in postsynaptic population. The population r receives the spread of activity across the matrix of synapses W multiplied by the presynaptic activity p (e.g. representing the aspects of the memory). The population r also receives additional direct postsynaptic activity q representing other individual events. Associative memory function requires that at least a subset of synapses have different external sources for presynaptic and postsynaptic input. Standard models of the hippocampal formation [4–6, 25, 57] focus on associations formed at modifiable synapses which are not the predominant influence on postsynaptic activity [58]. In these models, the direct external input driving postsynaptic activation would be synaptic input from the entorhinal cortex or dentate gyrus.

Equations 1 and 2 are the standard equations for associative memory in numerous models. However, these equations are unstable and sensitive to interference between memories [1, 13]. Most associative memory models avoid this instability and interference by ignoring synaptic transmission during Hebbian synaptic modification, but here we focus on instability as an important property. The instability can be seen by combining the equations by replacing r in equation 1 with r from equation 2, this results in the following equation:

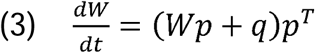

The instability of this equation is shown in Figure 2K-O. The figure shows that retrieval of a previous associative memory can influence the synaptic modification process during the encoding of a new overlapping associative memory. In this example, Figure 2K. shows the case of a new episode that involves a single point of overlap in the presynaptic input with a previously encoded memory (the place). For example, you meet a second colleague at the same place where you met the first colleague. The overlap results in spread across previously strengthened synapses (Thick blues lines, with the specific synapse involved in retrieval marked by the red line in Figure 2K). Thus, the same place can cause retrieval of the face from memory 1 (orange circle in Figure 2K). This causes strengthening of additional incorrect synapses from the place in memory 2 to the place in memory 1 as shown by the orange line in Fig. 2L. This problem becomes compounded further when learning another episode in succession in which there is one or more overlapping events with any previously encoded memories, as shown in Fig. 2P-T. This eventually results in a weight matrix W that cannot discern any previously encoded memories from one another with this problem resulting in unnecessary and undesired strengthening of many additional synapses.

The instability of the equation can be shown by solving equation 3 as a first order differential equation for patterns p and q to compute the change in synaptic strength over time. This yields a solution that grows linearly with presynaptic activity p and direct post-synaptic activity q, but also grows exponentially with presynaptic activity p:

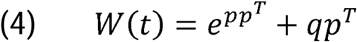

Note that this exponential growth concerns not only the growth in strength of individual synapses, but also the spread of undesired growth to synapses that should have no growth (orange lines weights in Figure 2L and 2Q) even if individual synapses are not allowed to grow beyond a certain maximum value. The exponential growth of synaptic weight in this equation demonstrates one major driving force that is proposed to underlie the pathology of early Alzheimer’s disease. This erroneous modification involves more than just excitotoxicity because it concerns increased synaptic modification at early stages rather than just increased neural activity. However, in some cases it could have the side-effect of excitotoxicity depending on how synaptic growth influences activity among particular subsets of neurons.

The solution to equation 3 can be made more stable by adding features that correspond to mechanisms that regulate biological synaptic transmission and synaptic modification. For example, as shown in Figure 3D, presynaptic inhibition of synaptic transmission during synaptic modification can prevent the exponential growth. This can provide stability in the equation by reducing the influence of prior weight W on synaptic modification. Abstract models of associative memory commonly avoid the instability using implementations of the equations that simply ignore retrieval during the encoding of new associations [7, 59][13,14]. Thus, stability of associative memory function could be obtained by ensuring that synaptic modification is combined with presynaptic inhibition of synaptic transmission during synaptic modification. Studies implementing optogenetic manipulations of LTP and LTD have demonstrated that the hippocampus can be programmed using simple Hebbian learning driven by LTP [60–62] resulting in the creation and ablation of synthetic memories in mice, so it is necessary to provide biological mechanisms that ensure that the induction of these functional synaptic modifications do not lead to the instability innate in Hebbian learning. Physiologically, this could be obtained by secretion of sAPPα causing an encoding phase that includes both enhancement of LTP (encouraging modification for encoding of memories) and presynaptic inhibition of synaptic transmission via presynaptic GABA_B_R1a receptors, whereas less secretion of sAPPalpha would correspond to the retrieval phase.

**Figure 3.**
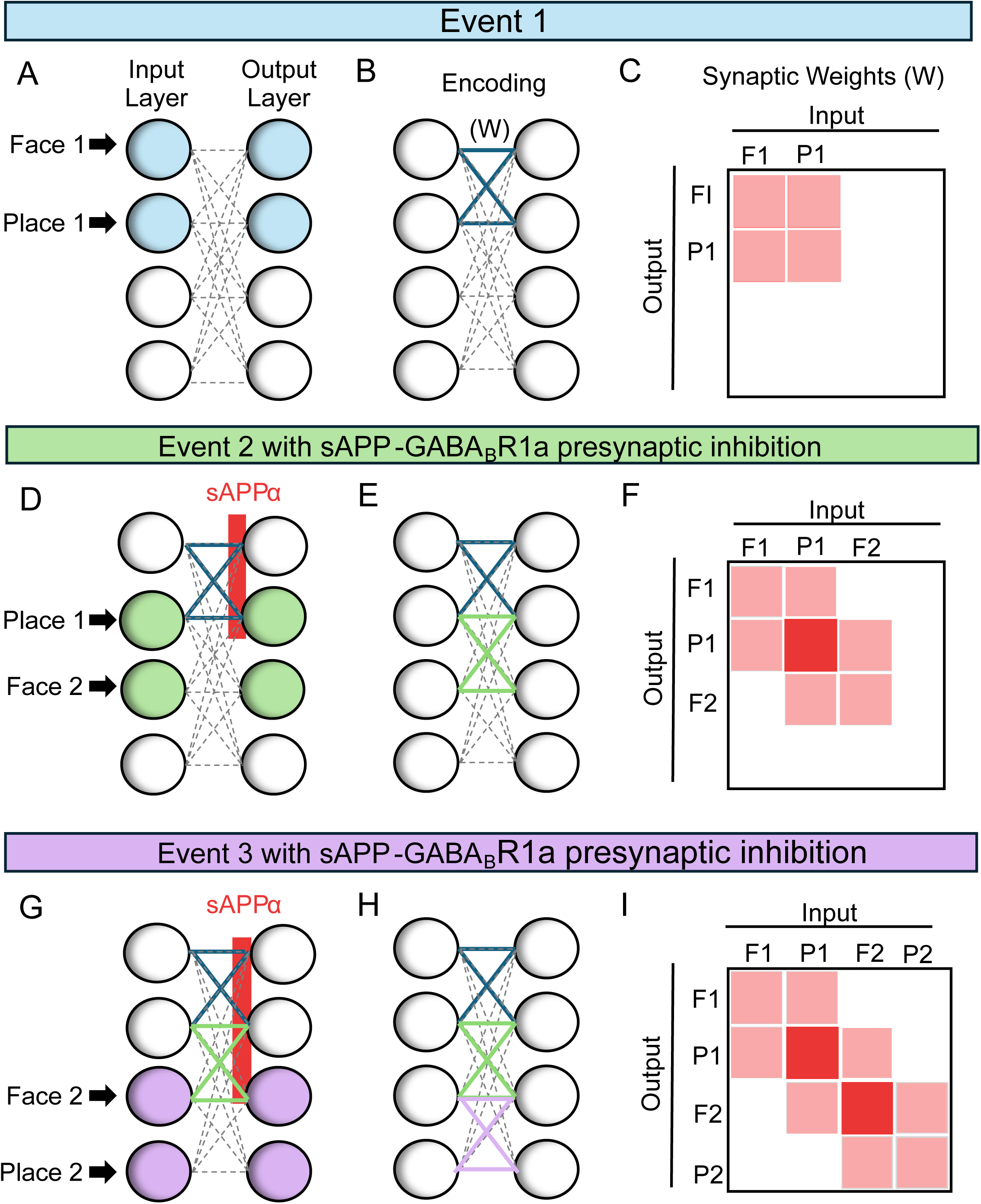
Presynaptic inhibition due to the activation of GABA_B_ Receptors by sAPPα prevents interference during encoding between overlapping associations. A) In the first event, one population of activated neurons (left blue circles) responds to a sensory input representing a face and a place associated with the memory, and this also activates the output population (right blue circles). B) During encoding, Hebbian synaptic modification strengthens synapses between the activated neurons (blue lines) and leads to an increase in sAPPα secretion at those synapses. C) The same pattern of strengthened synapses can be represented by pink shaded squares in a matrix of connections. D) Now, in a consecutive overlapping event 2 in which the activated neurons (green circles) have overlap in the same place being associated with a novel face, this overlap no longer causes undesired retrieval due to GABA_B_R-mediated presynaptic inhibition by sAPPα at the previously modified synapses. E-F) This results in Encoding LTP strengthening only desired new synapses (green in E, pink in F). G) Event 3 input only activates neurons receiving direct input (violet circles). Due to sAPP*α* presynaptic inhibition of prior retrieval, the Hebbian modification only strengthens desired synapses (violet lines in H, pink shaded squares in I).

Recently, the capacity of classical Hopfield networks has been increased by the use of modern Hopfield networks [63, 64]. These networks do not define a synaptic weight matrix as in classical Hopfield networks, but instead define an energy function and an associated activation function (update rule). Thus, it is difficult to directly map these new networks to actual biological synapses and synaptic modification rules. However, the energy functions in those models appear to implicitly define synaptic interactions as the sum of a function of the dot product of every stored pattern with the state pattern, which does not take into account their sequential dynamics of retrieval that would require the implementation of their update rule/activation function separately for each stored pattern. Thus, the new models make the same assumption of encoding as a separate process from retrieval as in the original Hopfield networks.

#### 2.1.2 Stabilizing the Hebbian learning equation

The stability of associative memory can be evaluated mathematically for conditions when encoding and retrieval are not assumed to be completely separated. This could be useful for understanding the interaction of disparate physiological effects that include presynaptic inhibition as well as the regulation of synaptic modification (both potentiation and depression). These effects involve different molecular pathways, but might all converge on the enzymatic processing of APP by both alpha- and beta-secretase pathways, so the equation could provide a unified vision of the functional role of APP processing on synaptic modification and network stability. The point is to demonstrate that sAPPα can cause effects that shift dynamics between encoding and retrieval, so that the network maintains better associative memory function. In this equation for stability of associative memory [1], the influence of molecular factors (*τ*) on synaptic modification, the influences (Z) on postsynaptic activity, and the effect of transmission across synapses W can be combined into a single equation:

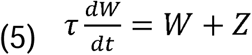

Using the standard solution for an inhomogeneous first order differential equation, this yields the solution:

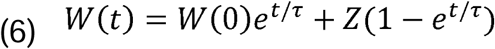

This corresponds to the solution for a standard nonhomogeneous first order equation [65] as described with the following structure and solution:

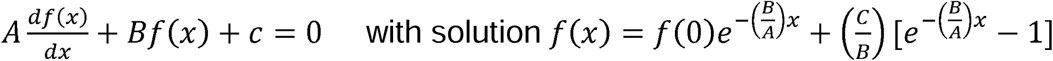

This equation can grow to a stable asymptote when 1/*τ* is negative or will show exponential growth when 1/*τ* is positive. The values from the previously presented equation 3 are Z=qp^T^, and 1/*τ*=pp^T^, but these components can be expanded to consider other molecular and physiological features influencing synaptic function. For example, equation 3 can be expanded to include a range of physiological parameters in the following form:

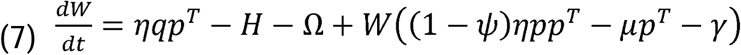

This equation includes parameters that are modulated by sAPP*α* or other modulators during an encoding phase but not during subsequent retrieval. These include the parameter *ψ* that reflects the magnitude of presynaptic inhibition of synaptic transmission during learning, *μ* for magnitude of synaptic decay dependent on presynaptic activity, *γ* for magnitude of synaptic decay that depends only on current strength, and *η* the learning rate that characterizes the magnitude of change dependent on presynaptic activity. H represents postsynaptic inhibition, and *Ω* represents a postsynaptic activity threshold for synaptic modification.

Separating out the components of this equation gives:

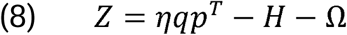

And

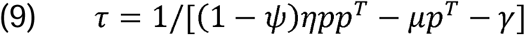

The 1/*τ* parameter describes the time course of exponential growth of synaptic strength. The exponential growth can be prevented if the parameter *ψ* reduces W to zero during encoding. For example, if sAPP-GABA_B_R1a presynaptic inhibition (which corresponds to *ψ* in the equation) causes sufficient presynaptic inhibition of synaptic transmission during encoding, this prevents exponential growth as shown in Figure 3. (A unique property of this mathematical analysis is that the presynaptic inhibition of synaptic transmission by GABA_B_R or m4 muscarinic receptors is mathematically equivalent to gated decay of synaptic strength based on both presynaptic and postsynaptic activity.)

The exponential growth can also be slowed by causing long-term depression based on current weights (corresponding to the decay of weight in proportion to the constants *γ* and *μ*). The learning rate parameter *η* could correspond to the role of sAPP*α* in forming synapses or enhancing synapse strength, whereas the A*β* generated by the beta-secretase pathway might regulate the synaptic decay parameters *γ* and *μ* involved in breaking down synapses.

Equation 9 predicts how mutations that affect physiological function of APP and APP secretion could influence the rate of progression of Alzheimer’s disease. Mutations that impair the presynaptic inhibition by sAPPα or the long-term depression due to A*β* should result in a more rapid time course of progression. Mutations that enhance presynaptic inhibition should result in a slower time course or absence of disease.

### 2.2 Large scale simulations

Simulations in MATLAB tested the basic associative memory function illustrated in Figures 2-3 using the expanded versions of equations 3 & 4, but with larger populations of neurons (i.e. larger vectors of presynaptic and postsynaptic activity) and multiple different vectors. In the simulations, each association involves external input that activates six neurons out of the 100-200 neurons in the presynaptic population of neurons to represent features of an episodic memory. These are associated with direct postsynaptic input that activates six neurons out of the 100-200 neurons in the postsynaptic population of neurons. In Figure 4, twenty new associations are stored, and each new input pattern overlaps with the previous association by one element.

**Figure 4.**
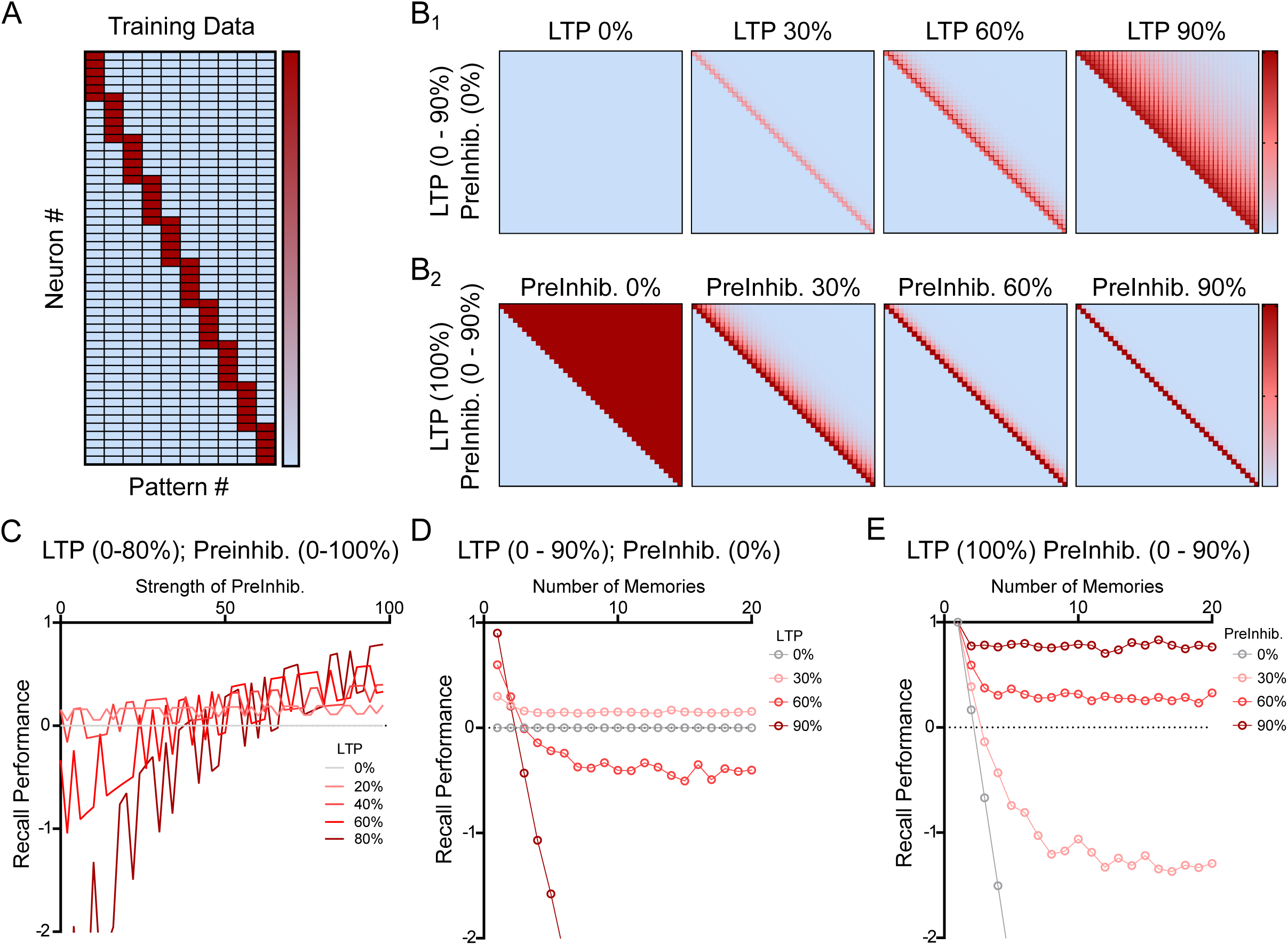
The combination of sAPPα mediated effects on both long-term potentiation and presynaptic inhibition stabilize Hebbian associative memory in simulation. A) Sample of memory training patterns generated to test the network recall accuracy of each iteration of the network. X axis corresponds to different patterns (pattern #) shown at different time points. Rows on the y axis indicate different neurons (neuron #) active in each input pattern (red indicates activity). B) These two rows show simulated weight matrices resulting after training on the full set of patterns with different parameters. Pink and red indicate strength of synaptic weights. The percent values show magnitude of effects of sAPPα on long term potentiation (LTP) and presynaptic inhibition (PreInhib) that were increased in strength to illustrate effects on runaway synaptic modification (RSM). B1). Weight matrices for 0% PreInhib, with LTP strength increased from 0% to 90%. At 0% LTP, no weight changes occur and no memories are stored. With 30% LTP only weak memories are stored (pink line along diagonal). With 60% LTP, weak memories are stored with interference from adjacent memories (broad line on diagonal). With 90% LTP strength and 0% PreInhib, the traditional Hebbian network dynamics lead to RSM, which strengthens synapses linking new memories to many previously presented memories (due to RSM, red squares representing synapses fill the upper right diagonal of the square matrix). B2) Matrices for LTP set at 100% with increases in sAPPα-mediated PreInhib from 0% to 90%. On the left, LTP at 100% and PreInhib at 0% results in massive RSM, maximally strengthening synapses between current memory and all previous memories (red values for all the undesired weights in upper right of square matrix). As %PreInhib increases to 30%, 60% and 90%, the retrieval of old memories during encoding is reduced, and the matrix of red synaptic connections contains less and less undesired synapses between the elements of each association (fewer red squares in upper right diagonal of square matrix). The combined effects of sAPPα on LTP and PreInhib stabilize the learning of the basic Hebbian network. C) Quantification of the recall performance for different values of increased LTP and PreInhib. On the left side with PreInhib of 0%, increased LTP causes interference that reduces Recall Performance to negative values. On right side, with PreInhib at 90%, as LTP increases, the performance gets better (reaching over 0.8) and gets closer to perfect recall (1.0). D) Quantification of memory retrieval performance for 0% PreInhib across increasing numbers of memories. 0% LTP results in no memory storage and 0% performance (gray circles). Performance is slightly better for low levels of LTP (30%, pink circles)), but drops off quickly as LTP increases further due to RSM (going to negative values for red circles for 60% LTP and dark red circles for 90% LTP). E) Performance for 100% LTP and PreInhib at 0% shows a sharp decrease in Recall Performance (gray circles). Performance gets better for larger values of PreInhib until showing best performance for PreInhib of 90% (dark red line above 0.8 performance for all memories).

The postsynaptic population activity is computed according to equation 2 to incorporate both the external input pattern and the retrieval that would occur in the network. The synapses are then strengthened according to equation 3. As an additional component, to show that runaway synaptic modification is due to interference causing a spread of synaptic weight through the network, rather than allowing unlimited strengthening of individual weights, the model was simulated with a limit on synaptic weights of 1.01. Despite this limit on weight, the network still exhibits runaway synaptic modification (RSM) as shown in the large-scale simulations.

## 3. RESULTS

### 3.1 sAPP*α* activation of presynaptic GABA_B_ receptors prevents runaway synaptic modification

Using the above-described methods, the phenomenon of runaway synaptic modification summarized in Figures 2-3 was simulated with sequential learning of a much larger number of overlapping associations, each with six input landmarks and six individual events. Figure 4A shows how each new input vector (pattern # on x axis) activates a sequential set of neurons (neuron # on y axis) with one overlapping element with the preceding input vector, similar to the simple example in Figure 2. The small and uniform number of inputs was selected to allow visualization. The simulations shown in Figure 4B-C model the biological components during encoding that include effects of sAPPα on synaptic modification [41, 42, 44, 45, 48, 49] and presynaptic inhibition of glutamatergic synaptic transmission by sAPPα [35], whereas they test performance in a retrieval phase with weaker effects of sAPP*α.* Data shows that LTP is enhanced by sAPPα which also simultaneously causes presynaptic inhibition of synaptic transmission via GABA_B_R. We first describe an introductory overview of the simulations in Figure 4, and we will then describe them in detail. Figure 4B1 shows that the enhancement of LTP can provide increases in encoding of individual memories, but this can also cause a breakdown in function termed runaway synaptic modification (RSM) for higher values of LTP (i.e. 90%). Figure 4B2 starts with LTP at 100% and shows that addition of stronger presynaptic inhibition prevents runaway synaptic modification even when LTP is strong. Thus, combining the two biological effects of sAPPa during encoding results in more stable memory function of the network as tested by performance with retrieval dynamics without sAPP*α*, as shown for 90% presynaptic inhibition and 100% LTP in the right-hand panel of Figure 4B2. After encoding, the matrix of synaptic connectivity shown in the right hand panel Figure 4B2 with higher levels of sAPP-GABA_B_R1a modulation strengthens the relevant memory synapses on the diagonal similar to what was shown in Figure 3F, which shows strengthening only along the diagonal of groups of four synapses between the elements of each association.

To describe the same effects in the figure in more detail, the row of synaptic matrices in Figure 4B1 shows how the function of the associative memory network breaks down for overlapping memories when Hebbian synaptic modification is applied at different strengths (0% to 90%) with the absence of the presynaptic inhibition of synaptic transmission (PreInhib=0%) during synaptic modification (Fig. 4B1). For LTP=0%, there is no strengthening and no memories are stored. This appears as an absence of red squares in the left panel of Figure 4B1. For LTP=30%, there is weak strengthening of synapses, shown as a pink diagonal. For LTP=60%, the network shows interference between memories that appears as a widening of the distribution of weights along the diagonal, indicating that each memory is interfering with the preceding memories. For LTP=90%, the build-up of interference due to retrieval of previous associations during learning of new associations causes runaway synaptic modification (RSM). This results in strengthening of many additional synapses that results in modification (red) of almost all of the synapses in the upper right section of the matrix. The matrix of connectivity has the same form as shown Figure 2T, showing that the strengthening of undesired synapses continues during subsequent associations to result in synaptic strengthening that links the elements of each new memory to almost every previously stored memory. Simply stated, the effect of interference during learning can be powerful and widespread, affecting an enormous number of additional synapses. This right panel shows bad memory retrieval performance as described later for Figure 4D.

In contrast, figure 4B2 shows that increases in presynaptic inhibition (PreInhib) can prevent runaway synaptic modification. The left panel of Figure 4B2 shows the synaptic matrix after learning with LTP at 100%, but PreInhib at 0%. This shows extreme RSM that modifies weights between every new memory and all the previously stored memories (solid red in upper right). The large difference in sum of synaptic weights with runaway synaptic modification is despite the fact that each individual weight in the matrix is only allowed to grow to 1.01 in the simulation. In contrast to the left panel, the subsequent panels show that even with LTP kept at 100%, larger amounts of presynaptic inhibition prevent this RSM, with weaker interference at PreInhib=30% and 60% and no interference or RSM at PreInhib=100%. This last panel on the right figure 4B2 shows that LTP=100% and PreInhib=90% allows strong memory weights (dark red along the diagonal) but prevents interference and RSM (no weights in upper right). This is similar to the example of how presynaptic inhibition prevents excess synaptic strengthening in Figure 3I. This right panel also corresponds to the best retrieval performance across memories shown in Fig 4E and described next.

The memory performance was tested by evaluating the accuracy of memory retrieval by the network when given only two random input cues for each memory and using feedback inhibition to set a threshold so that six postsynaptic neurons are activated each time during retrieval. The memory retrieval performance is shown on the y axis in Figure 4C for different levels of presynaptic inhibition (x axis) and for different strengths of LTP (gray to red curves). Similarly, the memory performance is shown on the y axis in Fig. 4D & E as the number of memories increases on the x axis. As an overview, Figure 4D shows performance when PreInhib=0% and LTP is increased to different values (gray=0% LTP and red=90% LTP). Figure 4E shows performance when LTP=100% and PreInhib is increased (gray=0% PreInhib, and red=90% PreInhib).

To provide more detailed description, Figure 4D tests performance for the matrices shown in Figure 4B1 and shows that when presynaptic inhibition is absent (PreInhib=0%), network function breaks down with higher values of synaptic modification (LTP). Setting the synaptic modification learning rate (LTP) to 0% results in no strengthening for any memory, and the memory retrieval performance remains at zero for all memories (gray line in Figure 4D). With an increase in LTP to 30%, there is some strengthening of synapses and memories are encoded weakly, resulting in performance that stays at a low level (pink line in Fig. 4D). For LTP=60%, there is more strengthening, but the strengthened synapses for each association causes more retrieval of previous memories during new encoding. As described in Figure 2, this results in interference that strengthens additional undesired synapses in the weight matrix (Figure 4B1) and during memory retrieval testing this results in retrieval of undesired components of previous patterns. This causes performance to fall below zero for more than three memories (red line for LTP=60%). Finally, for LTP=90%, the strong retrieval of previously stored memories during encoding causes extensive strengthening of undesired synapses forming associations with previous memories which causes runaway synaptic modification and quickly causes a rapid collapse of memory performance (dark red line that drops rapidly to negative values in Fig. 4D).

Figure 4E shows the performance for the matrices shown in Figure 4B2, in which the synaptic modification learning rate (LTP) was kept at 100% and we tested different values of presynaptic inhibition (PreInhib=0 to 90%). As a summary of this figure, the memory performance stabilizes when simulations were performed with addition of higher values of presynaptic inhibition of synaptic transmission by GABA_B_R1a receptors, as shown by the memory performance above 0.8 for PreInhib=90% in Fig. 4E. Figure 4E shows that with LTP=100% and presynaptic inhibition (PreInhib) at 0%, the performance drops rapidly (gray line in Figure 4E) because each memory cue retrieves interference from all previously stored memories, due to the undesired synaptic weights filling the upper right diagonal of weights in the connectivity matrix in the left panel in Figure 4B2. In contrast, encoding with presynaptic inhibition by sAPPα causes strengthening of only desired synapses in proportion to the number of stored memories and the number of additional synapses strengthened for each new memory (Fig. 4B2). As shown in Figure 4E, presynaptic inhibition by sAPPα-GABA_B_R1a with values of 60% to 90% prevents interference from prior retrieval during encoding. With PreInhib=60% the interference is weaker and performance increases to values above zero, as shown by the red line staying above zero in Figure 4E. With PreInhib=90%, there is no interference during learning and therefore the memory performance during retrieval shows better stability, with the dark red line showing performance above 0.8 for 20 stored memories. (Note that this function could continue for larger numbers of stored memories, and larger memory capacities can also be obtained with changes in other parameters including sparseness of network storage).

### 3.2 sAPP**α** is more effective at stabilizing associative memory than baclofen in associative memory models

The 1a isoform of GABA_B_R is predominantly localized to presynaptic compartments, where it inhibits glutamate release; whereas, the 1b isoform is predominantly localized to postsynaptic compartments, where it mediates postsynaptic inhibition [66]. While sAPPα specifically interacts with GABA_B_R1a, Baclofen is a non-specific agonist of both 1a and 1b-containing GABA_B_R heterodimers. Therefore, we sought to compare the effectiveness of sAPP and baclofen on stabilizing associative memory. We first confirmed that the properties of the GABA_B_R1a-dependent effects of sAPP on presynaptic inhibition was preserved in the expanded Hebbian equations (7 & 8). With high levels of LTP (100%), both encoding and memory recall performance are stabilized by sAPPα-mediated presynaptic inhibition at 60% (strong dark red line on diagonal of the right hand figure in Fig. 5A, similar to the right panel in Figure 4B2). This resulted in strong performance the dark red line over 80% in Fig. 5B (similar to Fig. 4E). We then tested effects of postsynaptic inhibition alone to model effects of R1b on the baseline instability of the Hebb learning dynamic (Fig. 5C). With low values of postsynaptic inhibition (0% or 30%), the underlying instability remained leading to runaway synaptic modification. For higher levels of postsynaptic inhibition (60%), the reduction of activity during encoding prevented RSM from occurring (right panel in Fig. 5C). However, this resulted in insufficient synaptic modification (weaker pink line on diagonal) and poor (near zero) memory recall performance (dark red line at zero in Fig 5D).

**Figure 5.**
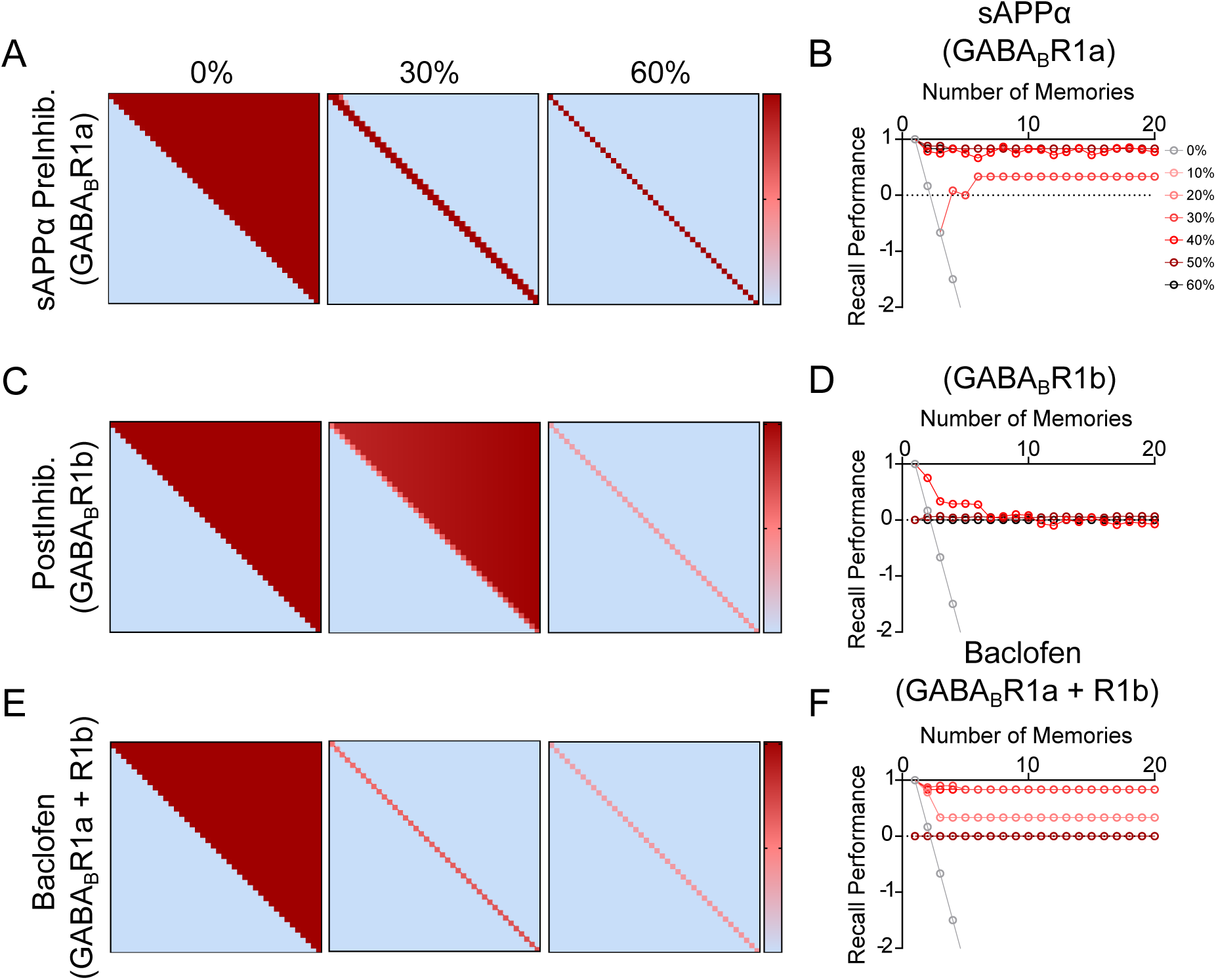
sAPPα is more effective at stabilizing associative memory than baclofen in associative memory models. A,C,E) Sample networks generated using an expanded Hebbian learning algorithm with LTP at 100% and different activation of subtypes of GABA_B_ receptors. A) Synaptic matrix shown after encoding with increasing levels (0%, 30%, 60%) of sAPPα-mediated activation of GABA_B_R1a. C) Matrix after encoding with increasing activation (0%, 30%, 60%) of GABA_B_R1b. E ) Matrix after encoding with baclofen-mediated activation of both GABA_B_R1a and GABA_B_R1b (at 0%, 30% or 60%). Each of these interventions stabilizes networks as shown by transition from many undesired strengthened synapses (red) in upper right of square matrix for 0%, to only strengthened synapses on the diagonal for 60%. However, performance is best for selective sAPPa activation of GABABR1a (A), in contrast withGABA_B_R1b (C) or baclofen-mediated activation of both GABA_B_R1a and GABA_B_R1b (E) in which the synapses at 60% are strengthened less (lighter pink) due to postsynaptic inhibition from GABA_B_R1b activation resulting in a weak association matrix (maximum value of 0.4) with a narrow window of baclofen efficacy. B,D,F). Quantification of memory performance over increasing number of memories (x axis) with different circles indicating increasing levels (0-80%) of sAPPα-mediated activation of (B) GABA_B_R1a, (D) activation of GABA_B_R1b (D), or (F) baclofen-mediated activation of both GABA_B_R1a and GABA_B_R1b. B) Memory Recall Performance was optimal (near 1.0) with sAPP-mediated activation of GABA_B_R1a (dark red circles near 1.0 in figure B). D). This contrasts with Recall Performance that is poor for only increased activation of GABA_B_R1b postsynaptic inhibition (dark red circles at 0.0 for most increasing number). F) This also contrasts with lower performance for baclofen-mediated activation of both GABA_B_R1a and GABA_B_R1b.

We modeled the effects of baclofen on both 1a and 1b-containing GABA_B_R heterodimers by modeling simultaneous changes in both presynaptic and postsynaptic inhibition using the weight matrix update functions described in equations 7,8, and 9 (Fig 5E). At weak pre and post synaptic inhibition (e.g. 30%), there is some weakening of the encoding of previous memories (Fig 5E, middle panel) but relatively stable recall performance (near 1) (Fig 5F, red line). Simulations with a combination of strong pre and post-synaptic inhibition (e.g. 60%) could prevent RSM (Fig. 5E, right panel), but this resulted in insufficient synaptic modification (pink line) and poor (near 0) memory recall performance (Fig 5F, dark red line for 60%, though note that lower levels have better performance). Together, these results suggest that sAPPα is more effective at stabilizing associative memory than baclofen and may help explain the limited success of baclofen as a treatment strategy for AD and other neurodegenerative diseases.

### 3.3 With random selection of memory activity, presynaptic inhibition by sAPPa-GABA_B_R1a signaling prevents runaway synaptic modification

For clarity of illustration, the simulations in the previous figures employ artificial patterns of memories, with each new memory of 6 elements containing one overlap with the previous memory and 5 new elements in numerical order. This was done as the simplest manner to illustrate the build-up of RSM. In Figure 6, the six presynaptic and postsynaptic active input elements are chosen randomly with a uniform distribution, so that the overlap and its frequency is random. RSM builds up in the same way if memories with 6 active elements are randomly added to the network, as shown in Figure 6A, which shows the same analysis for memories in which the 6 active elements are selected at random with random amounts of overlap across the entire bank of training data. Here we can again demonstrate the benefits of stabilizing Hebbian learning via presynaptic inhibition though sAPPα-GABA_B_R1a activity. Wherein high amounts of presynaptic inhibition by sAPPα-GABA_B_R1a result in improved encoding (Fig 6B) and memory performance (Fig. 6C).

**Figure 6.**
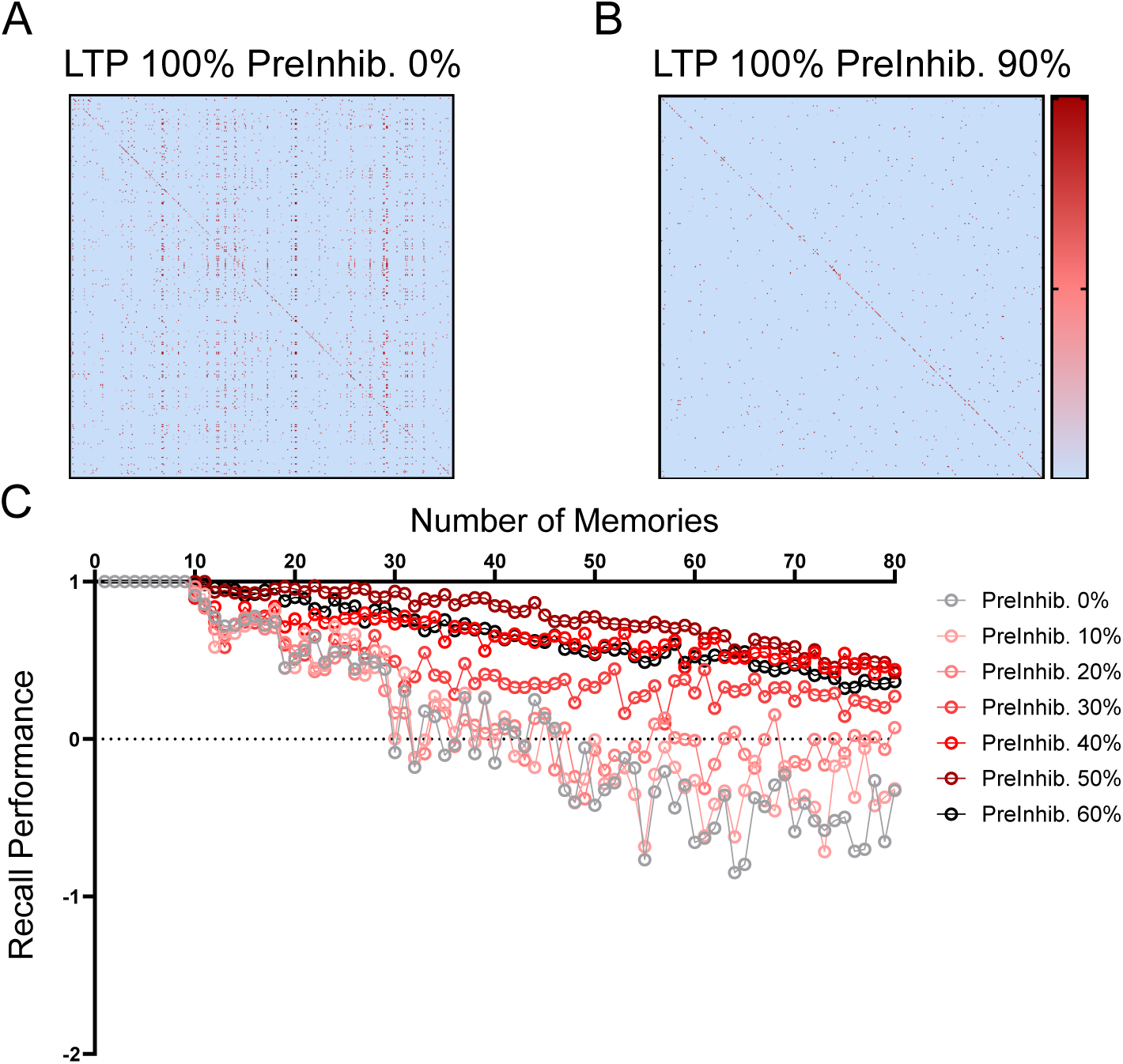
sAPPα prevents runaway synaptic modification in models of random memories. Larger sample networks (400X400 neurons) were generated to test effects of sAPPα-mediated activation of GABA_B_R1a with randomly distributed memory patterns and LTP at 100%. A) Without sAPPα-mediated PreInhib (0%), the 100% LTP induced synaptic modification that resulted in RSM (many small red squares that are mostly undesired synapses). B) Strong sAPPα-mediated PreInhib (90%) prevents RSM (reduction in non-specific synaptic growth). The synaptic matrix shows only desired synapses necessary for associative memory (fewer small red squares). C) Quantification of memory performance across sequential encoding of 80 random memories. Recall performance is strongest when sAPPα mediated presynaptic inhibition (PreInhib) is 50% (dark red circles), allowing excellent recall for a few stored memories and decaying gradually due to gradually increasing overlap of the random memories. In contrast, lower levels of sAPPα allow more interference during encoding leading to steep losses in memory function due to recall intrusions (e.g. gray circles at 0%).

### 3.4 sAPP***α*** prevents runaway synaptic modification without leading to insufficient modification

To summarize the effects of sAPPα and baclofen on synaptic modification in our models, we have categorized the sum of synaptic weights in the sample networks into three bins: stable (linear growth of synaptic weight and number), runaway (exponential growth of synaptic weight and number), and insufficient stable, and insufficient (static or too minimal growth of synaptic weight and number to achieve recall), across a range of tested strengths. Without presynaptic inhibition (PreInhib. = 0%), LTP (over a range of 0-100%) produces runaway synaptic modification in 100% of the strengths tested (Figure 7A, left).

**Figure 7.**
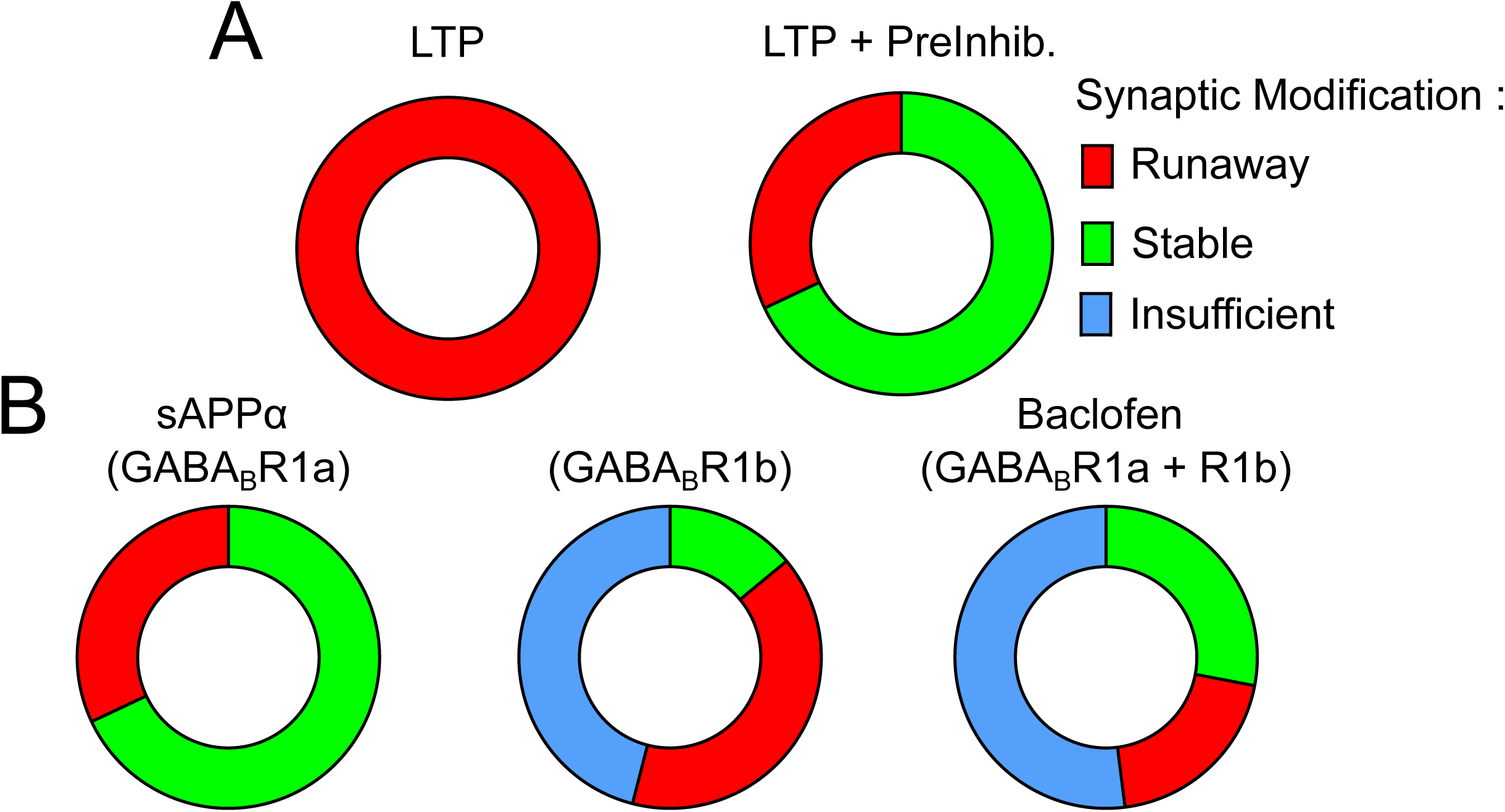
Summary of the protective role of sAPPα on network stability. The effects of synaptic inhibition are categorized into three unique bins based on the associated changes to the sum of synaptic weights of the sample network. Runaway synaptic modification (RSM, red) is defined by an exponential increase in synaptic weight per memory encoded. Stable (green) is defined by linear increase in synaptic weight per memory encoded. Insufficient (blue) is defined by an increase in weight insufficient to evoke recall. A (Left): The results of LTP (over a range of 0-100%) without presynaptic inhibition (Preinhibit. = 0%) gives RSM in all cases (red). A (Right): This contrasts with LTP (100%) with presynaptic inhibition (0-100%) which gives many conditions that show stable memory function (green). B (left): LTP at 100% with GABABR1a stimulation (Preinhib = 0-100%, Postinhib. = 0%) gives mostly stable memory function (green), B (middle): In contrast, LTP at 100% with GABABR1b stimulation alone (Preinhib = 0%, Postinhib. = 0-100%) gives mostly insufficient (blue) or RSM (red) and only some stable memory function (green). B (right): LTP at 100% with both GABABR1a and GABABR1b stimulation (Preinhib = 0-100%, Postinhib. = 0-100%) gives mostly insufficient memory (blue) with smaller amounts of stable memory function (green) and some RSM (red). These results are a summary of the total sum of synaptic weights across 100 sequential 40 memory simulations regarding each of the five parameters.

When GABA_B_R1a-sAPPα signaling is modeled (Preinhib. tested over a range of 0-100%) under conditions of high synaptic modification (LTP = 100%), runaway synaptic modification is observed in 24% of the tested strengths (red), with the remaining being stable (green, 68%) (Fig 7A, right; 7B left). If instead of presynaptic inhibition by GABA_B_R1a, postsynaptic inhibition by GABA_B_R1b is modeled under conditions of high synaptic modification (LTP = 100%, Preinhib. = 0%, PostInhib.= 0 -100%), similar percentages of runaway synaptic modifications are obtained (38%); however, only 18% of the strengths tested led to stable synaptic modification (green) with the emergence of insufficient synaptic modification (blue, 46%) (Fig. 7B, middle). When baclofen is modeled by combining activation of both GABA_B_R1a and GABA_B_R1b (LTP = 100%, Preinhib. = 0-100%, Postinhib. = 0-100%) (Fig. 7B right), stable synaptic modification was observed in 32% of the ranges tested (green), runaway in 22% (red), and insufficient in 46% (blue). Together, these results highlight that sAPPα and baclofen can both prevent runaway synaptic modification; however, baclofen does so at the expense of causing insufficient synaptic modification.

## 4. DISCUSSION

### 4.1 A protective role of sAPPα in Alzheimer’s disease

Our simulations demonstrate how sAPPα could be one of many possible mechanisms for preventing a proposed breakdown of synaptic homeostasis in Alzheimer’s disease. A protective role of sAPP is supported by experimental studies showing that sAPPα can improve cognitive outcomes in AD mouse models [67–69] . While human biomarker studies for sAPPα and sAPPβ have been variable in their directionality, they suggest that levels of sAPPα and sAPPβ may be dysregulated in AD [70–72]. Additionally, functional ADAM10 secretase levels are disturbed in AD [73] and point mutations in ADAM10 have been identified to increase AD susceptibility [74], further implicating changes to sAPPα in AD. GABA_B_R protein expression is altered in mouse models of AD further linking the synaptic effects of APP to severity and onset of disease [75].

Enhancing sAPPα has long been proposed as a strategy to protect against AD [69]; however, recent clinical trials have administered siRNAs that target APP as a strategy to reduce Aβ production to treat AD [37]. Thus, it is critical to consider potential consequences of targeting APP and its fragments in AD.

Figure 2-6 show how Hebbian associative memory function in standard models can be stabilized by the alpha-secretase production of sAPP*α* that causes presynaptic inhibition of glutamate release via activation of GABA_B_ receptors [35, 36] and also enhances synaptic modification (LTP) of glutamatergic synapses [41, 42]. This dual function of sAPP*α* prevents retrieval of previously encoded episodic memories from interfering with the encoding of new episodic memories. Simulations show that insufficient presynaptic inhibition allows runaway synaptic modification to spread through the network (e.g. Fig. 2T and Fig. 4B), which the malignant synaptic growth hypothesis [1–3] suggests is a functional cause of AD. Associative memories such as Hopfield networks have long been employed in modeling the hippocampus and other cortical structures [4, 57, 58, 76, 77] which informed our choice to use such a model [53, 78, 79].

These findings could be tested experimentally in AD mouse models by cannula injection of sAPPα or the short peptides composed of the binding region of APP to the GABA_B_R which mimics its function [35]. These studies could test for improved performance in one shot behavioral tests such as delayed non-match to position in a Morris water maze [80] or in other task designs that specifically include proactive interference [81, 82]. It is predicted that increasing sAPPα-GABA_B_R1a signaling will rescue or delay cognitive deficits in AD mouse models.

### 4.2 Runaway synaptic modification causes hyperactivation

Previous simulations of the model presented here indicate that runaway synaptic modification would be accompanied by hyperactivation in affected brain regions [1, 2, 12]. This prediction is supported by fMRI studies in humans with mild cognitive impairment showing increased activity in hippocampus during memory encoding [19, 83] and in young adults with the mutation that causes Alzheimer’s disease [20]. The hippocampal hyperactivation correlates with the appearance of amyloid plaques [84]. The hyperactivation also appears in cortical regions called the default network [85]. Runway synaptic modification could also contribute to the increased propensity for seizures observed in Alzheimer’s disease [86, 87].

Our modeling proposes that APP, by influencing both presynaptic inhibition and LTP, functions as a homeostatic mechanism consistent with a broader perspective on homeostatic plasticity in AD [88]. There are number of homeostatic synaptic mechanisms which contribute to the separation of encoding and retrieval by which existing modifications on different time frames contribute to conversion of novel stimuli into more robust semantic representations while not resulting in RSM [53, 54]. These mechanisms of homeostatic plasticity have been identified to converge on a number of shared molecular processes such as regulation of AMPA receptor recruitment to the postsynaptic density [89]. Hebbian plasticity may require mechanisms of Homeostatic plasticity for stability, [90] resulting a balanced system wherein perturbances to either homeostatic mechanisms or Hebbian mechanisms could result in unstable growth (RSM), or unreliable neural encoding [91]. It should be noted that certain activation states, like hyperactivity, limit the efficacy of homeostatic plasticity [92]. Signaling by sAPP is one candidate mechanism that may help regulate malignant synaptic growth in AD. Note that hyperactivation is here proposed to be caused by runaway synaptic modification but can also contribute to the excess synaptic modification. Previous papers about this model [2] have proposed that oligomeric A*β* inhibition of LTP [93–95] might be a homeostatic mechanism in which A*β* normally acts to reduce runaway synaptic modification, rather than A*β* being only a pathological neurotoxic mechanism. Our model confirms that reducing the learning rate *η* greatly slows the progression of runaway synaptic modification. However, it has also been shown that endogenously produced A*β* enhances evoked neurotransmitter release [96], which directly opposes the GABA_B_R-mediated effects of sAPP α on neurotransmitter release. Thus, sAPPα could be acting to counterbalance these effects of Aβ. The majority of the hyperactivation correlating with amyloid appearance has also been shown in Alzheimer model mice [97–99], but has been attributed to blockade of glutamate reuptake [100] rather than runaway synaptic modification.

### 4.3 Implications for pharmacological treatments

Modeling the role of presynaptic inhibition in preventing induction of AD suggests specific approaches for pharmacological treatment. Deficits in associative memory are among the first deficits identified among AD patient populations [101, 102]. Malignant synaptic growth would result in both retrograde and anterograde amnesia for recent episodic memories. Autobiographical memories integrate semantic memories and highly consolidated episodic memories that may have undergone more pattern separation that makes them more robust to interference, which could explain why they are lost at a later stage in AD. The therapeutic benefit of increasing presynaptic inhibition through molecular interactors like sAPPα in early AD would be a reduction or slowing of the progressive memory dysfunction.

Previously [2], it was proposed that AD could be slowed by enhancing selective presynaptic inhibition with agonists of the M4 muscarinic receptor [25, 58] . This could also be done via the presynaptic inhibition of glutamatergic synaptic transmission by presynaptically-located GABA_B_R [35], similar to muscarinic receptors [25, 58]. Presynaptic inhibition by GABA_B_R shows a selective effect on glutamatergic synaptic transmission at recurrent synapses but not afferent input synapses in the hippocampus [103, 104] and piriform cortex [105]. Presynaptic inhibition by GABA_B_R was previously proposed to prevent interference from prior retrieval during encoding based on enhanced LTP [26, 106] similar to the recently described dual effects of sAPP*α* that are modeled here.

Baclofen, an agonist of both 1a and 1b isoforms of GABA_B_R, has been proposed as a treatment for Alzheimer’s disease [107]. However, here we propose that specifically activating the 1a isoform of GABA_B_R, would more effectively reduce the progression of runaway synaptic modification in AD than a non-specific agonist. The discoveries that GABA_B_R1a is transported with APP [36] and activated by sAPP*α* [35] provides a target for the development of therapeutic strategies for modulating GABA_B_R1a-specific signaling. The identification of short (<17amino acid) peptides within sAPPα that are sufficient to bind GABA_B_R1a and mimic the effects of sAPPα on GABA_B_R1a provides a potential basis for the development of a GABA_B_R1a isoform–specific agonist.

Treatment could utilize activation of M4 muscarinic cholinergic receptors that cause presynaptic inhibition selectively at excitatory feedback glutamatergic synapses in the cortex [108, 109] whereas afferent input synapses show much weaker muscarinic presynaptic inhibition. The same synapses that show presynaptic inhibition also show strong Hebbian long-term potentiation in the hippocampus [25, 58] and the piriform cortex [110]. This is consistent with the anatomical distribution of M4 receptor labeling [111, 112]. Enhancement of presynaptic inhibition could utilize selective M4 receptor agonists and allosteric modulators [113, 114]. Modeling shows that the blockade of presynaptic inhibition by the muscarinic antagonist scopolamine should enhance proactive interference [5] and this was supported by behavioral data on effects of scopolamine in humans [115]. These previous models also showed how cholinergic modulation could be selectively activated for novel stimuli, based on a mismatch computed in region CA1 between current sensory input from entorhinal cortex and retrieval from region CA3. These papers were cited in a later paper that used the same mechanism for selective modification of synapses without synaptic transmission when there was a mismatch, allowing novelty-facilitated

Hebbian modification while blocking retrieval dynamics [116] . The APP protein could allow an additional type of novelty-facilitated modification, in which synapses that have not been recently modified (and therefore have more APP available for alpha-secretase splicing) might be more accessible for modification.

M4 receptor activators have been proposed as a treatment for hyperactivity observed in Alzheimer’s disease [117]. Treatment by selective M4 drugs for slowing progression would be most effective at an early stage of the disease before tangles have developed. Another possible pharmacological approach could address the role of group 1 metabotropic glutamate receptors in regulating long-term depression to counteract the mechanism of runaway synaptic modification. MgluR5, which is predominantly postsynaptically localized, has also been proposed as a therapeutic target in AD [118] . It possesses a similar kinetic profile as GABA_B_R but differs in the heterodimer assembly speed and surface retention [119].

Several characteristics of sAPPα position it as a particularly interesting candidate molecule to target therapeutically in AD. sAPPα as a shed extracellular domain would be expected to have slower turnover rates as a ligand at the synapse than classic neurotransmitters, such as GABA, glutamate, or acetycholine, sAPP is also of high therapeutic relevance because 1) it a 1a-isoform specific GABA_B_R ligand and 2) it is derived from the same precursor protein that leads to Aβ found in amyloid plaques associated with AD.

### 4.4 Summary

This model shows how cortical memory function could be stabilized by a secreted form of the amyloid precursor protein (sAPP*α*) that causes both presynaptic inhibition of glutamate release via activation of GABA_B_ receptors [35] and enhancement of long-term potentiation [42]. This joint function of sAPP can prevent runaway synaptic modification that could underlie AD. Moreover, modeling suggests that sAPP, which interacts with the 1a isoform of GABA_B_R, more effectively stabilizes associative memory than the non-specific agonist Baclofen. Together, these results support enhancing the sAPP-GABA_B_R1a signaling pathway as a potential therapeutic strategy for preventing cognitive impairment in AD. These findings also suggest caution in approaches that aim to reduce APP levels for the treatment of AD.

## Acknowledgements/Funding Sources

This work is supported by the National Institutes of Health, grant numbers R01 MH120073, R01 MH60013, R01 MH052090, R35 GM142726, and by the Office of Naval Research MURI N00014-16-1-2832 and MURI N00014-19-1-2571 and DURIP N00014-17-1-2304.

## Conflicts of Interest

H.C.R. is an inventor on a patent on the APP-GABABR interaction that is owned by VIB and the KU Leuven. Patent no. WO2018015296A1, “Therapeutic agents for neurological and psychiatric disorders.” D.B. and M.E.H do not have any conflicts of interest to disclose.

## Consent Statement

Human subjects were not used in this study.

## Notes

### Competing Interest Statement

The authors have declared no competing interest.

